# HUBMet: An integrative database and analytical platform for human blood metabolites and metabolite-protein associations

**DOI:** 10.1101/2025.09.16.676477

**Authors:** Xingyue Wang, Xiangyu Qiao, Alberto Zenere, Swapnali Barde, Jing Wang, Wen Zhong

## Abstract

Understanding human blood metabolites is essential for deciphering systemic physiology and disease mechanisms, yet remains challenging due to diverse origins and dynamic regulation. In this study, we developed HUBMet (https://hubmet.app.bio-it.tech/home), an open-access web server that includes 3,950 metabolites and 129,814 metabolite-protein associations, with four analytical modules: Over-Representation Analysis (ORA) for enrichment analysis; Metabolite Set Enrichment Analysis (MSEA) for quantitative data analysis; Tissue Specificity Analysis (TSA) for assessing metabolite-tissue relevance; Metabolite-Protein Network Analysis (MPNet) for identifying key metabolite-protein associations and functional modules. HUBMet’s utility is demonstrated through a COVID-19 case study revealing metabolic signatures associated with disease severity.

## Background

Blood metabolites are intermediate or end products of metabolic pathways that circulate throughout the human body. As a complete set of circulating metabolites, blood metabolomics can provide a systematic view of an individual’s health status, enabling the monitoring of physiological and pathological processes during disease progression [1,2]. Advancements in mass spectrometry (MS) and nuclear magnetic resonance (NMR) have significantly enhanced metabolomics, allowing for the identification and quantification of thousands of metabolites in human blood [3–5]. Efforts to catalog the blood metabolites, such as the Serum Metabolome Project and the Blood Exposome Database, have successfully identified extensive sets of endogenous and exogenous metabolites through high-throughput technologies and automated literature mining [6,7]. However, despite these advances, current human blood metabolite databases usually lack population diversity representation and show inconsistency across platforms. The size and composition of the blood metabolome vary significantly across studies, which affects the integration and downstream analysis of the human blood metabolome data. Moreover, analysis of the variability of blood metabolites remains challenging due to their diverse origins from various cells, tissues and organs [8,9]. Blood metabolite levels are highly sensitive to dietary and environmental changes, and individual metabolic variability further complicates analysis [10,11]. Thus, there is a pressing need for developing advanced bioinformatics tools and analytical platforms to interpret blood metabolite data and provide functional insights into the dynamics of metabolic processes in human health and diseases.

A variety of databases and analytical tools have been developed for metabolomic data analysis. For example, HMDB is a comprehensive source for chemical and biological annotation of metabolites [12], while tools such as MBROLE [13], WebGestalt [14], PathBank [15], and BioCyc [16] support metabolic functional and pathway enrichment analysis. Additional platforms like MetaboAnalyst, XCMS Online, and MMEASE [17–19], provide online tools for MS-based metabolomics data processing as well as downstream analysis including biomarker discovery, meta-analysis, and causal analysis. However, there remains a gap in tools specifically designed for the contextualized analysis of human blood metabolomics data, as most blood metabolites have multiple roles in human metabolism, which requires careful considerations of their tissue specificity and regulatory dynamics for functional interpretation.

Integrating blood proteomics with metabolomics provides a transformative approach to address these challenges. Metabolite-protein interactions, such as enzyme-substrate and transporter-ligand pairs, are central to metabolic regulation and cellular homeostasis [20]. Combining the comprehensive profiling of blood metabolites with proteomic data allows for a deeper understanding of the human metabolic network and its regulatory mechanisms. This integrative approach can also facilitate the discovery of novel metabolic pathways or functional modules through the analysis of metabolite-protein associations across different physiological and pathological conditions [21].

In this study, we developed HUBMet (HUman Blood Metabolites), an online web server for human blood metabolome analysis, built upon a curated database of human blood metabolites and metabolite-protein associations. HUBMet provides four analytical modules: (1) Metabolite Over-Representation Analysis (ORA); (2) Metabolite Set Enrichment Analysis (MSEA); (3) Tissue Specificity Analysis (TSA); and (4) Metabolite-Protein Network analysis (MPNet), to support functional enrichment analysis, tissue relevance assessment, and network exploration. The capability of the platform was further demonstrated through the analysis of a published COVID-19 metabolomics and proteomics dataset. This platform will contribute to advancing system-level analysis of blood metabolomics data and enhancing our understanding of metabolic regulation across health and diseases.

## Results

### Data summary and statistics of human blood metabolites

HUBMet contains 3,950 unique, detectable human blood metabolites identified through an extensive process combining literature mining and manual curation, representing data from 56 cohorts across 21 countries and regions (Fig. 1A-B, Additional file 1: Table S1-2). These metabolites were classified into nine categories, with the majority consisting of lipids (58.73%, n=2,320), followed by xenobiotics (18.33%, n=724), amino acid (9.65%, n=381), peptides (3.87%, n=153), carbohydrate (2.84%, n=112), nucleotide (2.40%, n=95), cofactors and vitamins (1.85%, n=73), energy (0.48%, n=19), and others (1.85%, n=73) (Fig. 1C). Notably, HUBMet includes numerous blood metabolites with reported clinical relevance that were not yet listed as blood metabolites in existing databases such as HMDB [12]. Examples include gamma-glutamylalanine, associated with chronic kidney disease incidence [22]; leucylalanine, elevated in colorectal cancer patients [23]; O-Ureido-D-serine, a potential treatment biomarker for Alzheimer’s disease [24]; and N6-acetyl-lysine, associated with aging [25].

**Fig. 1.**
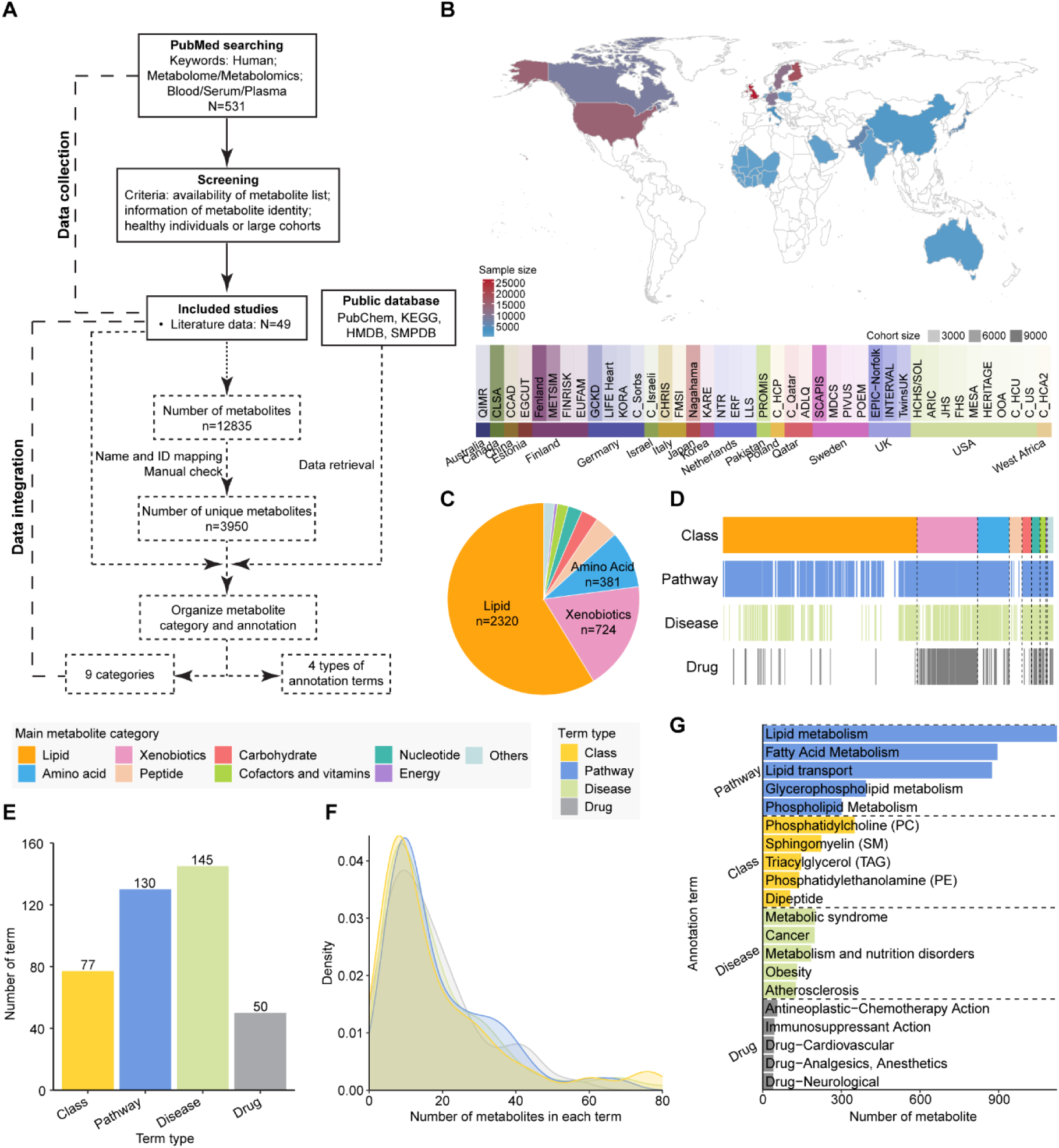
Human blood metabolite collection and annotation. **A** Workflow for blood metabolite data collection in HUBMet. **B** Top: World map showing the countries and regions covered, with colors representing total sample sizes per country or region. Bottom: Bar plot showing cohorts with more than 100 samples per country or region, colored by country/region, with transparency indicating cohort size. **C** Pie plot showing the proportions of metabolites across the nine metabolite categories. **D** Bar plot showing the different types of annotated terms (Class, Pathway, Disease, Drug) associated with each metabolite. **E** Bar plot showing the number of term sets, within each annotation type, that contain at least five metabolites. **F** Density plot showing the distribution of metabolites across term sets, with the x-axis representing the number of metabolites per term set (ranging from 0 to 80). **G** Bar plot showing the top five term sets with the largest number of metabolites in each annotation type. The x-axis indicates the number of metabolites in each term set.

Each of the 3,950 metabolites from the nine categories was classified into 162 chemical classes (Class) and annotated with the biological pathway (Pathway), associated disease (Disease), and the involvement in drug action and metabolism (Drug) (Fig. 1D). In total, 2,281 metabolites were linked to at least one biological pathway (Fig. 1D). Among xenobiotics, 304 out of 724 were involved in drug action and metabolism, while 803 metabolites were associated with various diseases. A total of 402 metabolite sets were created, each containing at least five metabolites (Fig. 1E). The number of annotation terms (class, pathway, drug, and disease) and the distribution of metabolites across these terms are shown in Fig. 1E and 1F. Most metabolite sets contained between 7 and 20 metabolites (Fig. 1F). The largest metabolite set, lipid metabolism, contained 1,122 metabolites (Fig. 1G), followed by related sets such as fatty acid metabolism, lipid transport, and glycerophospholipid metabolism. Key lipid classes in the human blood included phosphatidylcholine (PC), sphingomyelin (SM), triacylglycerol (TAG), and phosphatidylethanolamine (PE).

### Data summary and statistics of metabolite-protein associations

In total, HUBMet included 129,814 associations between 1,744 metabolites and 4,455 proteins, sourced from multiple approaches including curated database, human metabolic modeling, and statistical correlation analysis (Additional file 1: Table S3). The workflow for data collection is illustrated in Fig. 2A and Additional file 2: Fig. S1A, with a detailed procedure in Supplementary method (Additional file 2). Among all metabolite-protein associations in HUBMet, correlation analysis made the largest contribution, representing 56.1% (n=72,829) of the associations, involving 545 metabolites and 1,186 proteins (Additional file 2: Fig. S1A). Metabolic modeling contributed to 9.2% (n=11,905) of the associations, covering 516 blood metabolites and 2,643 proteins. Curated database from HMDB provided 37.48% (n=48,666) of the associations, covering 1,491 metabolites and 1,905 proteins [12]. To validate metabolite-protein associations, we utilized an independent cohort study and experimental data from the molecule-protein interaction database STITCH [26,27].

**Fig. 2.**
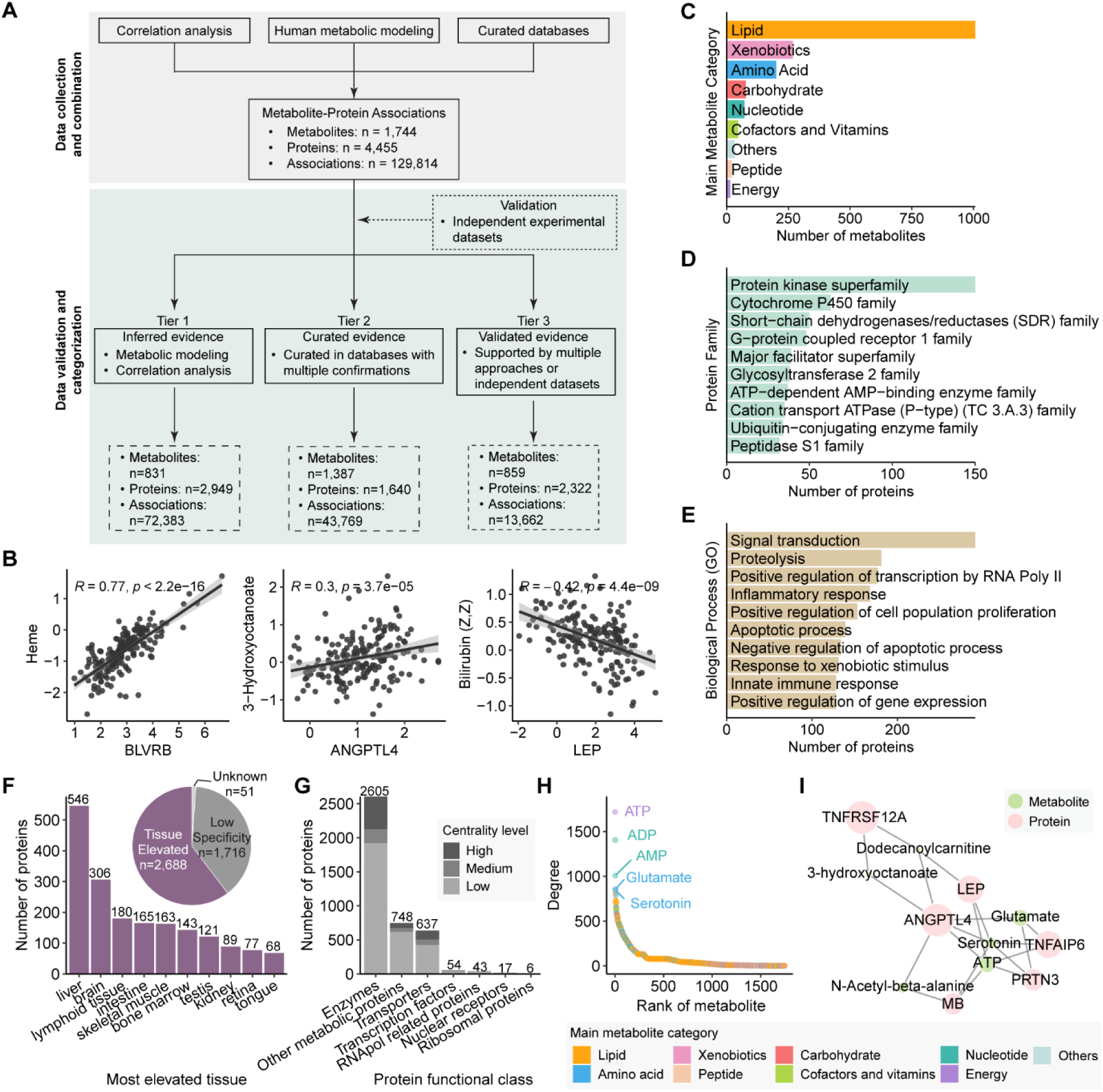
Data collection and construction of the metabolite-protein association network. **A** Overview of the workflow for the collection and integration of metabolite-protein association data. **B** Scatter plots showing correlation patterns for three selected metabolite-protein pairs in an independent validation dataset. Each plot includes the Spearman correlation coefficient (*R*) and *p*-value. Linear regression lines with 95% confidence intervals are displayed. **C-E** Bar plots showing (**C**) the main categories of the 1,744 metabolites with protein associations, and (**D**) the top associated protein families and (**E**) the major biological processes of the associated proteins. **F** Pie plot showing the tissue specificity of the associated proteins. Bar plot showing the top 10 tissues with the highest number of elevated proteins from HPA. **G** Bar plot showing the number of proteins in each functional class. The color code indicates the node centrality levels. **H** Dot plot showing the number of associated proteins (Degree) for each metabolite. The five metabolites with the most associated proteins are labeled. The color code indicates the metabolite categories. **I** Example subnetwork of metabolite-protein associations. Node size reflects the relative number of connections for both proteins and metabolites.

All associations in HUBMet were categorized into three tiers of evidence level based on the supporting evidence and validations: (i) inferred evidence, (ii) curated evidence, and (iii) validated evidence (details in Methods). In total, 72,383 associations were classified as Tier 1 (inferred evidence), 43,769 as Tier 2 (curated evidence), and 13,662 as Tier 3 (validated evidence) (Fig. 2A and Additional file 2: Fig. S1B). Examples of Tier 3 associations include BLVRB (Biliverdin reductase B, involved in heme catabolism) with heme, ANGPTL4 (Angiopoietin-like 4) with fatty acid 3-hydroxyoctanoate, and LEP (leptin) with bilirubin (Fig. 2B).

### Construction of metabolite-protein association network

The metabolite-protein association network was constructed based on the 129,814 metabolite-protein associations. The predominant classes of metabolites in the network include lipid (n=1,007), xenobiotics (n=269), and amino acids (n=202) (Fig. 2C). The metabolite-associated proteins belong to 1,092 families, with major families including the protein kinase superfamily (n=150), cytochrome P450 family (n=63), short-chain dehydrogenases/reductases (SDR) family (n=50) and G-protein coupled receptor 1 family (n=48) (Fig. 2D). Functionally, the most common biological processes related to metabolite-associated proteins include signal transduction (n=291), proteolysis (n=181), positive regulation of transcription by RNA polymerase II (n=177) and inflammatory response (n=168) (Fig. 2E). In terms of subcellular localization, the proteins in HUBMet were primarily located in cytoplasm (n=1,870), followed by cell membrane (n=1,085), nucleus (n=914) and secreted (n=815) (Additional file 2: Fig. S2A). Approximately 60% (n=2,688) of proteins in the network were classified as tissue elevated based on the annotation from the Human Protein Atlas (HPA) [28], with enrichment in liver, brain, and lymphoid tissue (Fig. 2F).

The centrality analysis of the metabolite-protein network revealed that proteins with high metabolite connections were predominantly enzymes, transporters, and other metabolic proteins (Fig. 2G and Additional file 2: Fig. S2B-C). Among them, monoacylglycerol O-acyltransferase 2 (MOGAT2), phospholipases (PLB1, PLCD3, PLA2G10, PLA2G1B), ATPase (ATP8A1, ATP10A, ATP8B1, ATP8B2), and apolipoprotein A5 (APOA5) were the top 10 proteins with the highest connectivity (Additional file 2: Fig. S2D). Notably, MOGAT2, a key enzyme involved in glycerol metabolism and enriched in liver and intestine, was associated with 42.7% (n=430) of lipids, including fatty acids, glycerophospholipids and glycerolipids, along with cofactors such as Coenzyme A (CoA) and Acetyl CoA, both of which play important roles in lipid metabolism (Additional file 2: Fig. S2D). In addition, key energy-carrying metabolites, including adenosine triphosphate (ATP), adenosine 5’-diphosphate (ADP), and adenosine monophosphate (AMP), showed extensive associations with proteins, mainly enzymes and other metabolic proteins (Fig. 2H and Additional file 2: Fig. S2E). For example, ATP was associated with numerous proteins involved in multiple biological processes, including ANGPTL4 and LEP (glucose homeostasis and insulin sensitivity), MB (oxygen transport and movement), PRTN3 (proteolysis and neutrophil extravasation), and TNFAIP6 (cell-cell and cell-matrix interactions during inflammation and tumorigenesis) (Fig. 2I).

### Community analysis of the metabolite-protein association network

To explore the clustering pattern of metabolite-protein associations, a Louvain method was employed for community detection [29], resulting in 12 distinct communities with varying numbers of metabolites and proteins (Fig. 3A-B). The enrichment analysis of Gene Ontology (GO) and KEGG pathways for each community provided further insights into their specific roles (Additional file 1: Table S4).

**Fig. 3.**
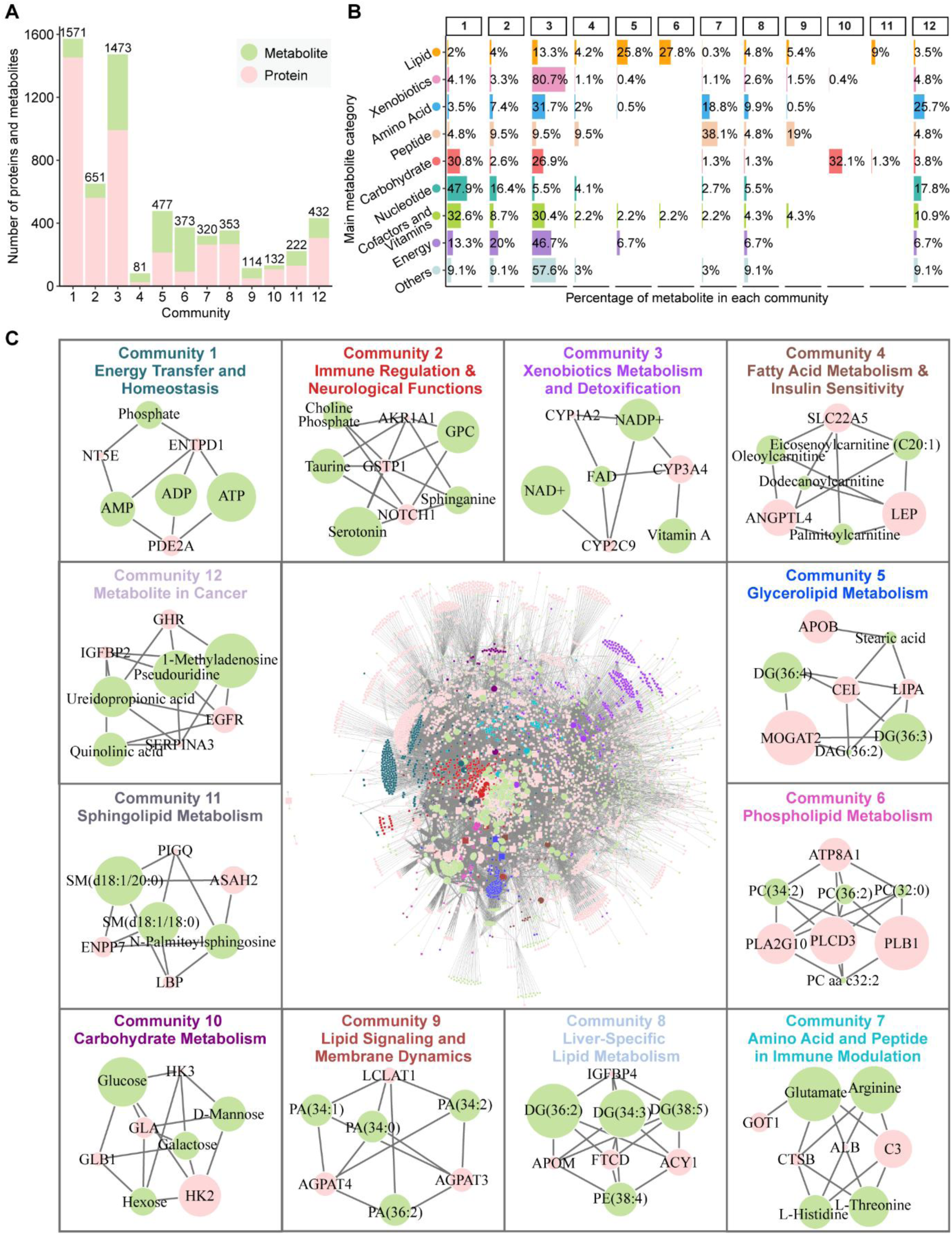
Community analysis of the metabolite-protein association network. **A** Bar plot showing the number of metabolites (green) and proteins (pink) in each community. **B** Bar plot illustrating the distribution of the nine main categories of metabolites across the twelve communities. **C** Overview of the metabolite-protein association network (inner plot), with individual community modules highlighted in different colors. Twelve subpanels display the hub proteins (pink) and metabolites (green) within each community. Node size represents the relative centrality of proteins and metabolites in each community.

The largest community, Community 1, was centered on energy transfer and homeostasis, comprising 1,453 proteins and 118 metabolites (Fig. 3C). Key components included energy transfer molecules ATP, ADP, and AMP. Enriched biological processes and pathways involved mitochondrial beta-oxidation, ubiquitin-mediated proteolysis, PPAR and AMPK signaling pathways, showing their significant role in energy metabolism and protein turnover. Community 2 was primarily associated with immune regulation and neurological functions, featuring key metabolites such as choline phosphate, serotonin, taurine, and glycerophosphorylcholine (GPC), along with hub proteins like NOTCH1 and GSTP1. Enrichment analysis highlighted pathways related to cytokine signaling, Notch signaling, and neuromodulation. The presence of serotonin, taurine, and GPC further showed the crosstalk between immune and neurological systems, suggesting overlapping roles in immune and brain function regulation (Fig. 3C). Community 3 involved a significant proportion of xenobiotics (80.7%, n = 217) and cofactors (30.4%, n=14) for xenobiotics metabolism and detoxification (Fig. 3B). Cytochrome P450 enzymes (e.g., CYP3A4 and CYP2C9) were major hub proteins, reflecting the community’s role in metabolizing drugs and detoxifying foreign compounds.

Fatty acid and other lipid metabolisms were represented across several communities (Fig. 3C). Community 4 emphasized fatty acid metabolism and insulin regulation, involving metabolites like oleoylcarnitine and palmitoylcarnitine, and proteins such as ANGPTL4 and LEP [30]. Community 5 was centered on glycerolipid metabolism, featuring diacylglycerol (DAG) and enzymes like MOGAT2. Community 6 involved phospholipid metabolism, highlighting structural lipids such as phosphatidylcholine (PC) and enzymes like PLB1, critical for membrane integrity and signaling [31]. Community 8 centered on liver-specific lipid metabolism, with liver-enriched proteins (e.g., ACY1, FTCD, APOM) and glycerophospholipids [32], emphasizing the liver’s central role in lipid regulation. Community 11 focused on sphingolipid metabolism, including sphingomyelin and ceramide catabolism, which are critical for cell signaling and apoptosis [33]. Key enzymes such as ASAH2 and ENPP7 were central to sphingolipid turnover [34].

Community 7 centered on amino acid and peptide metabolism with a focus on immune modulation (Fig. 3C). Key metabolites such as arginine and threonine were involved in immune responses and nutrient signaling [35–37], while complement component C3 served as a hub protein connecting amino acid metabolism to immune functions. Community 9 was associated with lysophosphatidic acid (LPA) metabolism, significant in lipid signaling and membrane dynamics [38], featuring enzymes like LCLAT1 involved in phospholipid remodeling. Community 10 centered on carbohydrate metabolism, incorporating key enzymes such as HK2 and GLB1 that were essential for glycolysis, gluconeogenesis, and carbohydrate catabolism [39], emphasizing its role in energy production. Community 12 was uniquely associated with cancer-related pathways, involving nucleotides and modified nucleosides like 1-methyladenosine, which were associated with multiple types of cancers [40]. Key proteins such as EGFR, a well-known oncogenic driver, were prominent, underscoring their role in oncogenesis and as potential therapeutic targets in cancers such as non-small cell lung cancer (NSCLC), breast cancer, and colorectal cancer [41].

### Tissue specificity analysis of blood metabolites

To investigate the relevance of blood metabolites across different human tissues, we conducted a tissue specificity analysis for blood metabolites in HUBMet. This analysis mapped 1,744 metabolites to 36 tissues and organs by integrating metabolite-protein associations from HUBMet with tissue-specific gene expression data from HPA [28] (Additional file 1: Table S5). Three criteria assessing the statistical overrepresentation of associated proteins in tissues were developed, and three types of tissue relevance were defined, including “Tissue Relevant”, “Low tissue relevance”, and “Unknown” (details in Methods). Based on these criteria, along with external validation from a curated database (HMDB), the reliability of metabolite-tissue relevance was further stratified into high, medium, and low reliability (Additional file 2: Fig. S3A-C, details in Methods).

As an example, hyodeoxycholate, a bile acid, was identified as liver-relevant with high reliability, supported by 11 of its 15 associated proteins, including CYP3A4 and UGT2B4, which were both enriched and highly expressed in the liver (Fig. 4A). Although hyodeoxycholate also had associated proteins elevated in the intestine and kidney, none met the criteria for tissue relevance. Another example was L-Tryptophan, a dietary amino acid, which was found to be relevant to multiple tissues and organs, including kidney, placenta, liver, intestine, parathyroid gland, lymphoid tissue, and bone marrow, with 26 out of the 50 associated proteins showing enrichment in these tissues (Fig. 4B). In particular, L-tryptophan was identified as a kidney- and placenta-relevant metabolite with high reliability, consistent with previous reports that tryptophan metabolism was associated with kidney function and diseases [42,43], and crucial for pregnancy and fetal development [44,45].

**Fig. 4.**
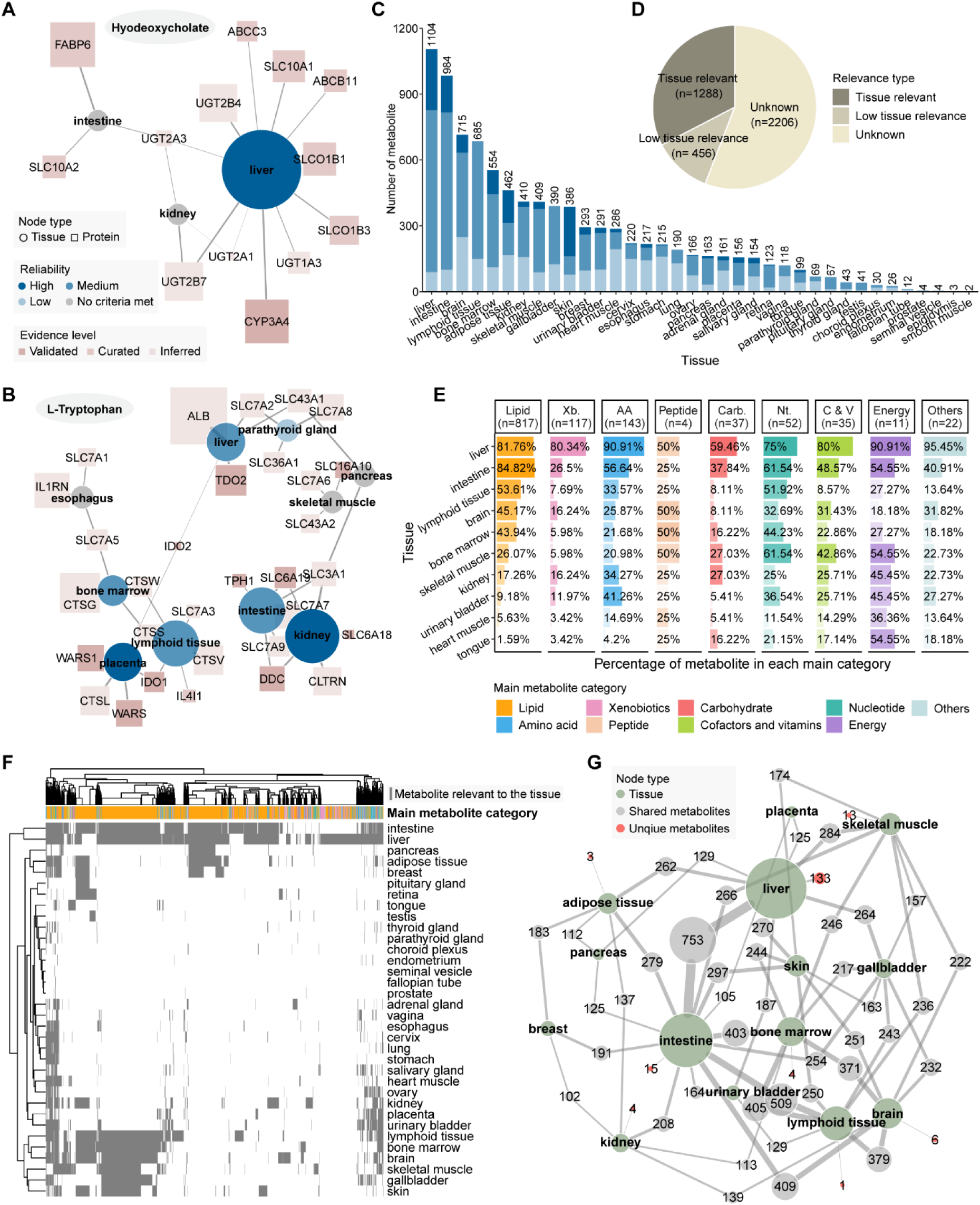
Tissue-specific relevance analysis of blood metabolites. **A-B** Tissue-elevated proteins associated with (**A**) Hyodeoxycholate and (**B**) L-Tryptophan. Node shapes represent protein (square) and tissue (circle). Tissue node size represents the relative number of associated elevated proteins in the tissue, with node color indicating the reliability level of tissue-specific expression. Protein node size represents the relative medium RNA expression across all elevated tissues, and the node color indicates the evidence level of the metabolite-protein association. **C** Bar plot showing the number of metabolites relevant to each tissue, stratified by three reliability levels. **D** Pie chart showing the distribution of metabolites across different tissue relevance annotations. **E** Bar plot showing the percentage of metabolites from each of the nine categories associated with the top ten tissues. These tissues were selected as the union of the top five tissues with the largest number of relevant metabolites in each metabolite category. **F** Heatmap showing the overlap of tissue-relevant metabolites across tissues. Each grey line represents a single metabolite. **G** Metabolite-tissue network showing the number of shared metabolites between each pair of tissues. Tissue node size represents the number of relevant metabolites in each tissue, while edge labels indicate the number of shared metabolites between tissue pairs. E-G include only metabolite-tissue pairs with a high or medium reliability level. Abbreviations: Xb., Xenobiotics; AA, Amino acids; Carb., Carbohydrate; Nt., Nucleotides; C & V, Cofactors and Vitamins.

In total, 1,288 metabolites were identified as “Tissue Relevant”, involving 9,252 metabolite-tissue pairs (Fig. 4C-D). Of these, 1,263 metabolite-tissue pairs had high reliability, 5,127 had medium reliability, and 2,862 had low reliability (Additional file 2: Fig. S3C). In addition, 456 metabolites were classified as “Low tissue relevance”, and 2,206 as “Unknown” (Fig. 4D). Metabolites with high or medium reliability of tissue-relevance were mainly enriched in tissues and organs involved in a broad spectrum of metabolic pathways, such as the liver and intestine (Fig. 4C). Among the 36 tissues analyzed, the liver contained the highest number of relevant metabolites with high and medium reliability (n=1,014, 81.9%), showing its central role in human metabolism (Additional file 2: Fig. S3D). Among the liver-relevant metabolites, 65.9% were lipids (n=668) (Fig. 4E). The liver also showed enrichment for 90.9% (n=130) of amino acids, including alanine, a key substrate for gluconeogenesis, and betaine, a methyl donor essential for homocysteine detoxification and methionine metabolism [46,47]. Furthermore, the liver was enriched for the largest proportions of xenobiotics (80.3%, n=94) and cofactors and vitamins (80%, n=28), including paraxanthine, ethyl glucuronide, vitamin A, NAD^+^, and bilirubin (Fig. 4E), aligning with the critical role of liver in detoxification and biosynthesis [48]. Energy metabolism-related metabolites, including fumarate, citric acid and succinic acid, which were essential components of the tricarboxylic acid (TCA) cycle, were also found to be liver-relevant.

Several other tissues showed specific enrichment of tissue-relevant metabolites, including immune-related organs such as lymphoid tissue and bone marrow, which were enriched in nucleotides (57.7%, n=30), such as adenosine, a mediator of immune response [49] (Fig. 4E). In addition, 14 nucleotides, including N2, N2-Dimethylguanosine and ADP, were specifically associated with genes elevated in the salivary gland (Additional file 2: Fig. S3D). Carbohydrates like glucose, galactose, and lactate were largely relevant to gene expressions in the liver (59.46%, n=22), intestine (37.84%, n=14), kidney and skeletal muscle (27.03%, n=10) (Fig. 4E). Amino acids were mainly relevant to the liver, intestine, urinary bladder, and kidney (Fig. 4E). Furthermore, two peptides, carnosine and leucylglycine, previously reported to be associated with brain cell activity modulation and physiological signaling, were found to be brain-relevant metabolites [50]. Tissues and organs with high metabolic activity, including liver, intestine, brain, lymphoid tissue, and bone marrow, shared a large proportion of tissue-relevant metabolites (n=288), the majority of which were lipids (n=259) (Fig. 4F). In particular, the liver and intestine shared the largest number of metabolites (n=753) (Fig. 4G). Among them, 278 phospholipids, which are key components of lipoproteins involved in the lipid transport between the liver and intestine, were identified as tissue-relevant metabolites for liver and intestine.

### Functional modules in HUBMet

HUBMet provides a suite of tools to facilitate the analysis of human blood metabolites (Fig. 5A). Each metabolite in HUBMet is accessible on its dedicated webpage with detailed annotations, including the chemical classification, biological pathways, involvement in drug action and metabolism, and disease associations (Fig. 5B). An interactive network of associated metabolites and proteins for each metabolite is presented, along with the functional annotations and tissue specificity of the associated proteins (Fig. 5B). By default, the network displays the top 10 proteins with highest centrality scores and the top 5 metabolites with highest Jaccard similarity scores (Fig. 5B). Users can expand the network by including additional associated proteins and metabolites. To ensure broad compatibility with existing metabolomics platforms, HUBMet supports 17 widely used metabolite identifiers, including Name/Synonym, HMDB, KEGG, BioCyc, PubChem, PDB, ChEBI, DrugBank, Phenol-Explorer, FooDB, KNApSAcK, Chemspider, VMH, LipidMaps, SwissLipids, SMILES, and InChI Key. Users can search for individual metabolites and run the analytical modules using any of these identifier types.

**Fig. 5.**
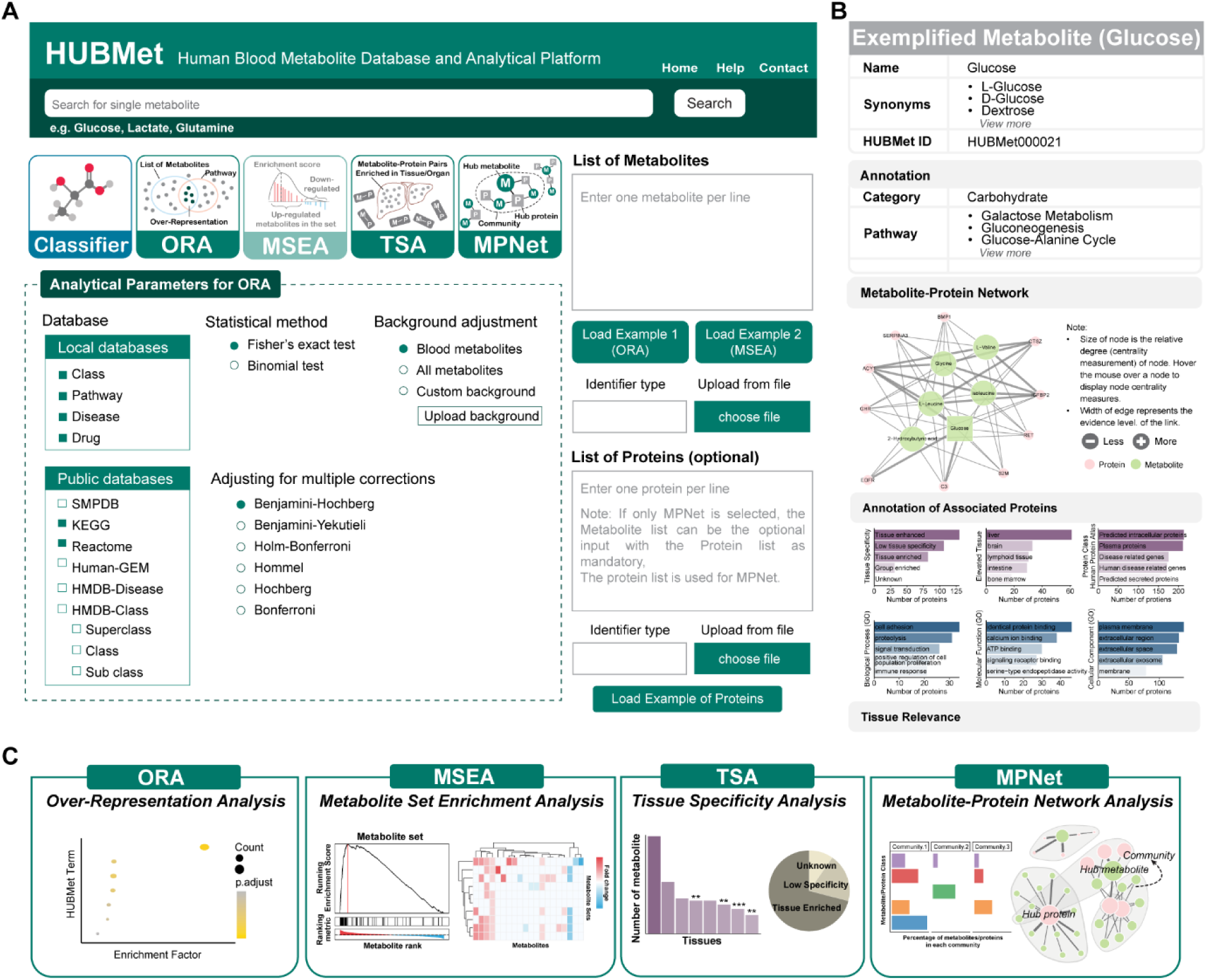
User interface and functional modules of the HUBMet Web Server. **A** The HUBMet homepage interface. **B** Example webpage of an example metabolite (glucose), showing the structured results from HUBMet, including basic information, annotations, associated metabolite-protein network, and relevant tissues. **C** Example visualizations from each of the four analytical modules, including (i) dot plot showing enriched terms from ORA; (ii) enrichment score plot and heatmap from MSEA; (iii) bar plot and pie plot summarizing tissue relevant metabolites from TSA; and (iv) community analysis and interactive network exploration from MPNet.

The HUBMet platform provides four analytical modules, including: (1) Metabolite Over-Representation Analysis (ORA); (2) Metabolite Set Enrichment Analysis (MSEA); (3) Tissue Specificity Analysis (TSA); (4) Metabolite-Protein Network Analysis (MPNet). Specifically, the ORA module employs two statistical enrichment methods, including Fisher’s exact test and binomial test, along with multiple test correction, to determine whether a predefined set of metabolites is significantly overrepresented in a user-specified input list. A default reference background of 3,950 human blood metabolites is available in HUBMet for the ORA analysis. The MSEA module applies methods adapted from gene set enrichment analysis (GSEA) to detect coordinated metabolic changes based on quantitative data [51–53], such as fold changes or correlation values. The MSEA method utilizes permutation testing and multiple hypothesis correction to calculate enrichment scores (ES), normalized ES, nominal *p*-values, and false discovery rates (*FDR*). Both ORA and MSEA utilize the 402 curated metabolite term sets in HUBMet categorized by class, pathway, drug, and disease annotations. In addition, the ORA and MSEA modules can interface with external public databases including KEGG, SMPDB, Reactome, Human-GEM, and HMDB [12,54–57] (Fig. 5A). The TSA module in HUBMet supports both tissue-relevance annotation and tissue enrichment analysis for user-provided metabolites. Each input metabolite is mapped to curated metabolite-tissue relevance in HUBMet. Enrichment analysis is then performed using Fisher’s exact test or binomial test, with multiple testing correction, based on predefined tissue-specific metabolite sets, enabling users to systematically identify tissues that are significantly enriched for the input metabolites (Fig. 5C). The MPNet module provides interactive exploration of metabolite-protein associations (Fig. 5C). Users can input lists of metabolites, proteins, or both, and filter associations by evidence level. By default, all background associations curated in HUBMet are included. In the resulting network, edges represent metabolite-protein associations, and centrality metrics, including degree, closeness, and betweenness, are computed to identify key network components. Furthermore, community detection via modularity optimization is applied to identify clusters of closely connected metabolites and proteins.

A step-by-step tutorial is available for download from the HUBMet web server (https://hubmet.app.bio-it.tech/help), which provides detailed guidance on the functional modules, supported input data types for each module, and parameter settings. In brief, the ORA module takes a user-defined metabolite list as input, with configurable parameters including the annotation database, statistical method, multiple testing adjustment approach, and background adjustment method. The MSEA module, in contrast, requires a complete list of identified metabolites along with their quantitative data (in a two-column format), and provides options for database selection and filtering based on metabolites set size. The TSA module requires a metabolite list as input, whereas the MPNet module accepts both metabolite list and protein list (optional). Supported metabolite identifiers are as described above, and supported protein identifiers include gene symbols, Ensembl IDs, UniProt IDs, and HMDB Protein IDs (HMDBP). All modules provide interactive visualizations and downloadable result files. Additionally, users can explore the web server using the provided example datasets and examine both interactive outputs and downloadable results.

### Case study

To demonstrate the analytical capabilities of HUBMet, we performed a case study using the plasma metabolome (n=1,018) and proteome (n=1,463) data from a COVID-19 study cohort [26]. The dataset included samples from controls (n=182), outpatients (n=183) and several/critical cases (n=272). Differential expression analysis identified 103 metabolites significantly changed in the outpatient group and 414 metabolites in the severe/critical group compared to the control group (Fig. 6A-B, Additional file 2: Fig. S4A, Additional file 1: Table S6).

**Fig. 6.**
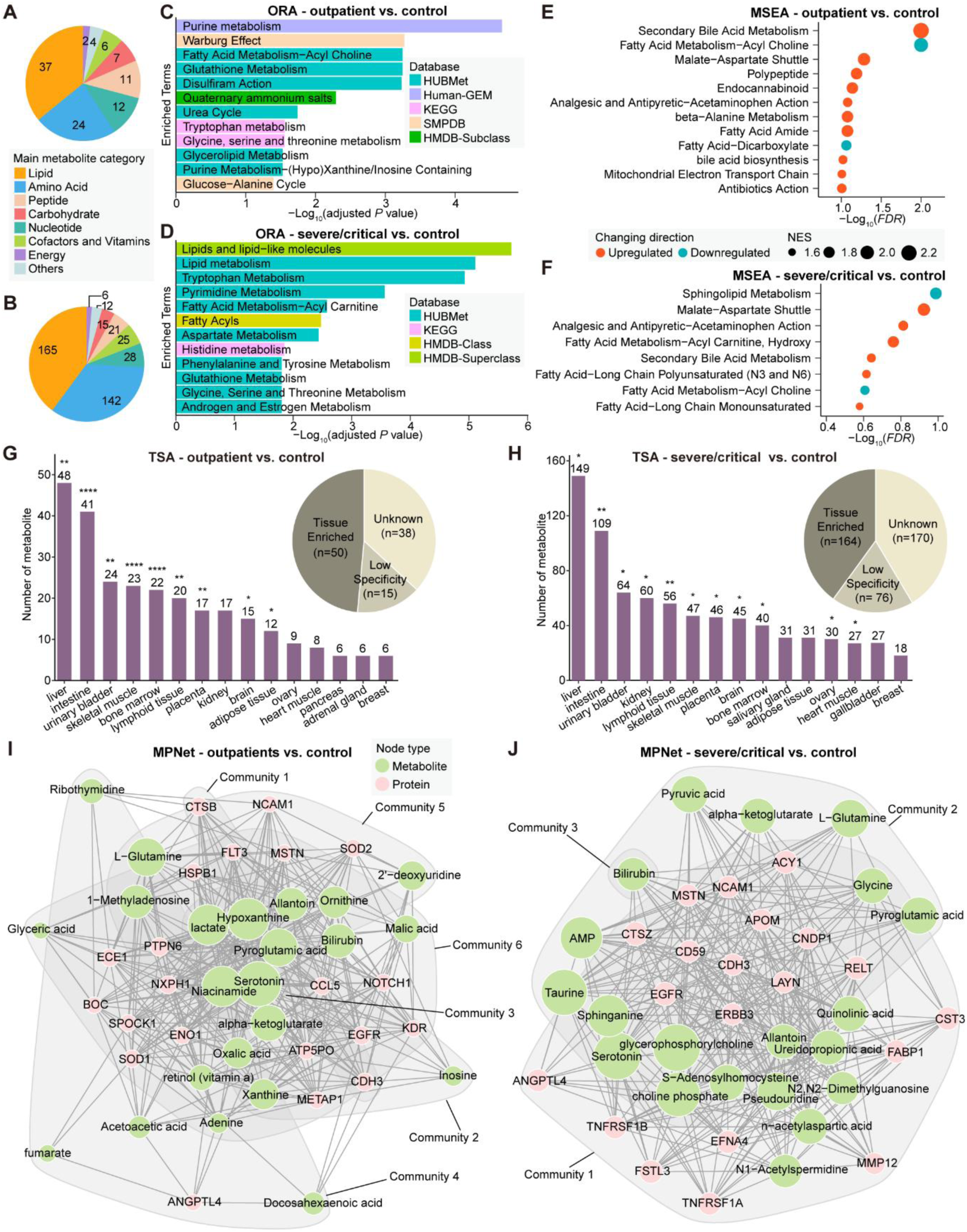
Comprehensive analysis of clinical COVID-19 metabolomic profiles using HUBMet. **A-B** Pie charts showing the main categories of differential metabolites identified in (**A**) outpatients and (**B**) severe/critical patients compared to controls. **C-D** Bar plots showing the ORA results for differential metabolites of (**C**) outpatients and (**D**) severe/critical patients compared to controls, with the x-axis as -Log10 adjusted *p*-value (Fisher’s exact test with Benjamini-Hochberg correction). **E-F** Dot plots showing the MSEA results for (**E**) outpatients and (**F**) severe/critical patients compared to controls, with the x-axis as -Log10 *FDR*. Dot size indicates the normalized enrichment score (NES), and the color shows the direction of changes in each metabolite set. **G-H** Bar plots and pie plots showing the TSA results for differential metabolites with tissue relevance in (**G**) outpatients and (**H**) severe/critical patients compared to controls. Pie plots show the tissue-relevant types of differential metabolites, and bar plots show the enriched tissues with the number of relevant metabolites. Significance levels are indicated by asterisks. **I-J** Metabolite-protein association networks derived from MPNet for (**I**) outpatients and (**J**) severe/critical patients, based on differential metabolites and proteins. Node size indicates the relative degree of proteins (pink) and metabolites (green) in the network. *adjusted *P* <0.05; **adjusted *P* <0.01; ***adjusted *P* <0.0001; ****adjusted *P* <0.0001

Using HUBMet, we performed a comprehensive analysis of differential metabolites, with the results from HUBMet in Additional file 1: Table S6. ORA revealed that outpatients exhibited significant dysregulation in key metabolic pathways, such as purine metabolism, Warburg effect, fatty acid metabolism (acyl choline), and glutathione metabolism (Fig. 6C). These alterations were consistent with previous findings and were likely associated with virus replication and inflammatory response [58,59]. In severe/critical cases, the main metabolic changes involved lipid metabolism, tryptophan metabolism, and pyrimidine metabolism, aligning with reported metabolic signatures associated with the severity of COVID-19 [60,61] (Fig. 6D). The MSEA results were consistent with these findings, showing upregulation of malate-aspartate shuttle and downregulation of fatty acid metabolism (acyl choline) (Fig. 6E). In addition, secondary bile acid metabolism was significantly up-regulated in both outpatients and severe/critical cases, supporting previous reports that elevated levels of bile acids were correlated with the severity of COVID-19 [62,63] (Fig. 6F). TSA results revealed that in both outpatients and severe/critical cases, metabolites showed significant enrichment in the liver and intestine, followed by the urinary bladder, skeletal muscle, and lymphoid tissue (Fig. 6G-H). These findings highlight the involvement of hepatic, gastrointestinal, and immune-related tissues in COVID-19, consistent with the systemic and multi-organ effects of COVID-19 [64].

To further investigate the molecular interplay between dysregulated metabolites and proteins, we applied the MPNet module, which integrated the metabolite-protein changes utilizing the curated metabolite-protein association network in HUBMet (Fig. 6I-J, Additional file 2: Fig. S4A-B). In outpatients, six communities were identified in the metabolite-protein network, corresponding to pathways involved in cell adhesion (community 1), energy and purine metabolism (community 2), serotonergic regulation (community 3), lipid metabolism (community 4), amino acid metabolism (community 5), and host-virus interactions (community 6). In contrast, the severe/critical patient group showed a more condensed network, comprising three major communities associated with lipid metabolism and immune signaling (community 1), nervous system and mitochondrial energy metabolism (community 2), as well as heme catabolism and amino acid metabolism (community 3). Interestingly, serotonin, a classic neurotransmitter, was identified as one of the hub metabolites in both networks and associated with proteins and metabolites related to immune and digestive systems (Additional file 2: Fig. S4C-D). This was consistent with previous findings that altered tryptophan metabolism, which was enriched in both ORA results, could lead to the decrease of serotonin and contribute to symptoms related to long-COVID [65]. Other shared hub metabolites identified in both networks included L-glutamine, pyroglutamic acid, alpha-ketoglutarate, bilirubin and allantoin, which were closely associated with SARS-CoV-2 replication, oxidative stress, immune dysfunction, coagulopathy and liver injury [66–69]. Comparative analysis of these two networks also revealed severity-related metabolites. In outpatients, central metabolites such as hypoxanthine, niacinamide (nicotinamide) and lactate were identified and associated with purine metabolism, immunity and energy metabolism, as well as aerobic glycolytic metabolism [70–73]. In contrast, severe/critical cases were characterized by key metabolites like glycerophosphorylcholine, sphinganine, and choline phosphate, which have been reported to be associated with inflammatory responses and served as biomarkers of COVID-19 severity [74–76].

## Discussion

Analyzing human blood metabolomics data poses significant challenges due to the complexity and heterogeneity of the human blood metabolome. Circulating metabolites reflect a dynamic interplay of endogenous metabolic activities and exogenous exposures, with origins from diverse tissues and organs [1]. While existing bioinformatics tools have advanced the processing and interpretation of metabolomics data, most remain fragmented and often lack context-specific functionality, particularly in the domain of blood metabolite interpretation. To address these limitations, in this study, we developed HUBMet, an integrative multi-omics analytical platform that enables in-depth functional exploration of human blood metabolome by integrating curated metabolite data with proteomic associations and providing streamlined analytical pipelines for metabolomics functional analysis.

In total, HUBMet curated 3,950 unique blood metabolites with annotations of biochemical classes, biological pathways, disease associations, and involvement in drug action and metabolism. Combining data from curated databases, genome-scale metabolic models, and population-scale correlation analyses, we constructed a comprehensive blood metabolite-protein association network, comprising 129,814 interactions that connected 1,744 blood metabolites with 4,455 proteins. Based on the constructed database, HUBMet provides four analytical modules, including: (1) Metabolite Over-Representation Analysis (ORA), (2) Metabolite Set Enrichment Analysis (MSEA), (3) Tissue Specificity Analysis (TSA), and (4) Metabolite-Protein Network Analysis (MPNet). The platform supports 17 different metabolite identifier types and includes analytical terms from external databases such as KEGG, Reactome, and SMPDB, ensuring broad compatibility.

A unique aspect of HUBMet is the integration of metabolite data with protein associations, which facilitates the biologically contextualized interpretation of blood metabolomics data, as metabolites and proteins usually function as interdependent molecular partners in various cell processes [21]. While existing tools can provide effective statistical analysis for metabolomics data, they typically do not incorporate multi-omics information, thereby limiting their ability to elucidate the mechanistic roles of metabolites in physiological and pathological processes. Based on the constructed metabolite-protein association network, HUBMet links the metabolite changes to enzymatic activity, transport functions, and broader regulatory pathways. Through the network analysis, we identified 12 distinct metabolite-protein communities, each enriched for specific biological functions, including energy metabolism, immune signaling, lipid processing, and neurotransmission. These findings revealed the diverse roles of circulating metabolites and highlighted the utility of the metabolite-protein association network in advancing system-level understanding of metabolic regulation.

Another advancement in HUBMet is the implementation of a tissue-specificity relevance analysis for blood metabolites. By leveraging tissue-specific gene expression data from the Human Protein Atlas and combining it with metabolite-protein associations, we established a computational framework to infer the likely tissue/organ origins or functional contexts of blood metabolites. In total, our analysis identified over 1,238 metabolites with high or medium confidence tissue relevance, providing a powerful resource enabling the users to interpret metabolite changes in relation to specific tissues or organs.

Furthermore, to support functional interpretation of metabolomic data, HUBMet implemented two complementary enrichment analysis methods, including ORA and MSEA. ORA was designed for the identification of significantly enriched annotation terms among input metabolite lists, while MSEA could detect coordinated changes across predefined metabolite sets using a modified GSEA framework. To reduce redundancy and improve interpretability, HUBMet applied a hierarchical clustering method to group semantically similar annotation terms across multiple databases and prioritized representative terms by statistical significance. This approach streamlined enrichment results and supported clearer biological interpretations of the analyzed metabolites.

We demonstrated the capability of HUBMet through a case study analyzing plasma metabolomic and proteomic data from a COVID-19 cohort [26]. The analytical results from the HUBMet web server provided new insights into the metabolic alterations after SARS-CoV-2 infection. HUBMet not only captured known metabolic perturbations, such as disruptions in purine metabolism, bile acid metabolism, and tryptophan pathways, but also provided novel insights into tissue-level involvement, implicating liver, intestine, and lymphoid tissues in disease pathology. Furthermore, MPNet analysis revealed key metabolite-protein hubs, such as serotonin, L-glutamine, and bilirubin, consistent with reported roles in immune response, inflammation, and COVID-19 severity [66,68,69,77]. These findings highlight the potential of HUBMet to advance the metabolite-protein-based biomarker discovery and provide novel mechanistic insights.

Despite the strengths, HUBMet has several limitations. First, although it represents one of the most comprehensive curated collections of blood metabolites from multiple populations to date, the database may still underrepresent newly discovered or low-abundance metabolites, especially as analytical technologies evolve. Second, the 56 included cohorts were predominantly European and largely comprised generally healthy individuals, which may limit the coverage of non-European populations and disease conditions, potentially reducing the applicability and analytical power in these contexts. Third, while many metabolite-protein associations were supported by multi-source evidence, comprehensive experimental validation remains a critical next step to ensure confidence for translational applications. Fourth, the tissue-specific relevance annotations of the curated blood metabolites based on gene expression profiles of the associated proteins, which, although informative and capable of providing valuable functional context, require further validation through direct tissue-specific metabolomic studies, which is currently a challenging task by the limited availability of large-scale tissue metabolomics datasets and variability in sample processing and quantification protocols. To address these limitations, future updates to HUBMet will include the integration of metabolites from more diverse populations and disease-specific cohorts, as well as low-abundance metabolites when high-quality datasets become available, and will expand the inclusion of experimentally validated metabolite-protein associations. In addition, as large-scale tissue metabolomics datasets become available, we aim to incorporate more direct evidence for tissue-specific annotations.

## Conclusions

HUBMet provides an open-access, user-friendly web server for the integrative functional analysis of human blood metabolome data. By combining curated metabolite annotations with protein associations and tissue-specific information, HUBMet enables systematic investigation of blood metabolomics through modular enrichment tools, tissue-specificity relevance analysis, and large-scale metabolite-protein network exploration. This platform will serve as a valuable resource to support metabolic biomarker discovery and the investigation of molecular mechanisms underlying human health and disease.

## Methods

### Data collection of human blood metabolites

A systematic search for publications on human blood metabolites over the past 10 years (up to August 20, 2024) was performed via PubMed using the following search terms: ‘blood’, ‘serum’, ‘plasma’, ‘human’, ‘metabolite’, ‘metabolome’, and ‘metabolomics’. In total, 531 publications were identified and reviewed. Exclusion criteria were applied to studies lacking an available list of detected human blood metabolites or insufficient identification information. Research focused on specific diseases or blood cells was also excluded. After applying these exclusion criteria, 49 publications were included for the construction of the list of human blood metabolites [11,21,58,78–123] (Additional file 1: Table S1).

The identified human blood metabolites were then mapped to HMDB ID (version 5), followed by careful manual curation. This resulted in a final list of 3,950 unique metabolites with 12,835 synonyms. Data from HMDB, SMPDB, KEGG, Human-GEM (version 1.18.0) and Reactome (accessed July 2023) were used for metabolite annotation [12,54–57]. The metabolites were classified into nine main categories, including Lipid, Xenobiotics, Amino acid, Peptide, Carbohydrate, Nucleotide, Cofactors and vitamins, Energy, and Others. Each metabolite was further annotated with (1) chemical characteristics (Class), (2) participation in biological pathways (Pathway), (3) associations with specific diseases (Disease), and (4) involvement in drug action and metabolism (Drug). Class and Disease annotations were obtained from the Chemical Taxonomy and Associated Disorders and Diseases sections of HMDB [12]. Pathway information was integrated from SMPDB, KEGG, Human-GEM, and Reactome [54–57].

Drug-related annotations were derived from SMPDB pathways categorized as Drug Action and Drug Metabolism [54].

### Blood metabolite-protein associations

Blood metabolite-protein associations were derived through an integration of three complementary data sources: (1) curated database (HMDB, version 5.0) [124], (2) human metabolic modelling (Human-GEM) [56], and (3) statistical correlations from two large-scale metabolomics and proteomics studies [21,125]. We further validated these metabolite-protein associations using blood metabolome and proteome data from 182 control plasma samples in the Mayo Clinic Biobank (USA) [26,126] and chemical-protein interactions in the STITCH (Search Tool for Interacting Chemicals) database [27,127]. The detailed data collection and validation process is provided in Additional file 2: Supplementary method.

Based on the supporting evidence and validation results, we developed a three-tier classification system to evaluate the confidence level of each metabolite-protein association: (i) Tier 1 (inferred evidence): associations derived from computational modeling or statistical correlation analysis; (ii) Tier 2 (curated evidence): associations obtained from manually curated databases; (iii) Tier 3 (validated evidence): associations supported by at least two different approaches or validated using independent datasets.

### Metabolite-protein network construction and analysis

An unweighted and undirected network based on all the metabolite-protein associations was constructed by using the R package igraph (v2.0.3) [128]. The centrality of each node (proteins and metabolites) was evaluated using the degree, closeness, and betweenness functions in igraph. Hub nodes were identified as those with the highest degree and closeness value (top 5) in the network. The centrality levels of proteins in the network were defined with degree score in log2 scale: degree score > median + 2×median absolute deviations (MAD) were classified as High, those with a score > median + 1× MAD and ≤ median + 2×MAD as Medium, and the remaining as Low. Jaccard similarity scores were calculated using the similarity function. Community detection was performed by using Louvain algorithm with a resolution set as 1.5. The subnetwork consisting of hub nodes was visualized in the study.

On the web server, the greedy optimization of modularity method (via cluster_fast_greedy function of igraph) is used for community detection to improve computational efficiency. For network visualization, the number of edges is limited to 100, with priority given to hub metabolites and proteins. Nodes belonging to the same community are highlighted with a grey background. Users can interactively update the network based on the selected evidence level of associations.

### Metabolite tissue relevance analysis

To infer the tissue relevance of blood metabolites, we integrated the gene expression data from 36 tissues in the Human Protein Atlas (HPA) with metabolite-protein associations curated in HUBMet [28]. For each metabolite, the set of associated proteins was analyzed across tissues using three complementary criteria to capture tissue-specific expression patterns. Criterion 1-Significant tissue-specific protein enrichment: For each tissue, the proteins annotated as tissue enriched, tissue enhanced, and group enriched in HPA were considered as tissue elevated. Fisher’s exact test was used to test if these tissue-elevated proteins were significantly overrepresented among the proteins associated with the metabolite. A *p*-value less than 0.05 was considered statistically significant. Criterion 2-High tissue-specific protein count: For each metabolite, we calculated the number of associated tissue-elevated proteins in each tissue. The distribution of these protein counts across all 36 tissues was used to compute a 95% confidence interval (CI), and a tissue was considered relevant under this criterion if its count exceeded the upper bound of the CI. Criterion 3-High tissue-specific protein expression: Protein-coding genes in HPA with normalized transcript per million (nTPM) higher than 1 were considered as detected. For each tissue, a Wilcoxon test was used to evaluate if the expression levels of the metabolite-associated proteins were significantly higher than the median expressions in all other tissues. A *p*-value less than 0.05 was considered as significantly elevated expression. Moreover, to avoid spurious associations, tissues with fewer than three tissue-elevated, metabolite-associated proteins were excluded from the evaluation.

Based on these criteria, each metabolite was assigned to one of the three tissue relevance categories. Metabolites meeting at least one of the three criteria were classified as “Tissue Relevant”; those failing all criteria were classified as “Low tissue relevance”; and those with insufficient information on protein associations or tissue-specific expression were labeled as “Unknown”. Furthermore, “Tissue Relevant” annotations were further stratified into three reliability levels: (1) High reliability: all three criteria met; (2) Medium reliability: Two criteria met, or one criterion met with supporting localization data from HMDB; (3) Low reliability: only one criterion met, without independent support. Only metabolite-tissue annotations with medium or high reliability were incorporated into the TSA module on the HUBMet web server.

### Over-representation analysis

Over-representation analysis (ORA) was implemented in the HUBMet web server to identify enriched functional terms among input metabolite sets. Users can select either Fisher’s exact test or the binomial test as the statistical method, both of which were implemented using the fisher.test() and binom.test() functions from the R stats package (v4.3.1) [129].

Annotation terms included metabolite classifications across four categories (Class, Pathway, Disease, Drug). To ensure statistical robustness, annotation terms with fewer than five metabolites were excluded. After filtering, 402 annotation terms remained available in HUBMet for enrichment analysis.

### Metabolite set enrichment analysis

The statistical framework for metabolite set enrichment analysis (MSEA) in HUBMet was adapted from the gene set enrichment analysis (GSEA) method [51–53]. The procedure involved three main steps, including enrichment score (ES) calculation, permutation testing, and multiple hypothesis testing correction. Detailed methods are provided in the Additional file 2: Supplementary method.

### Non-redundant enrichment term analysis

To reduce redundancy and improve the interpretability of the enrichment results, pairwise similarity between annotation terms was quantified using Kappa scores, which reflect the extent of overlap based on shared metabolites [130]. Enriched terms were first ranked by statistical significance (adjusted *p*-value in ORA or *FDR* in MSEA). Representative terms were then selected iteratively from the top of the list, ensuring that no pair of representative terms had a Kappa score greater than 0.3.

The remaining non-representative terms were then assigned to hierarchical clusters by associating each with the most statistically significant representative term to which it has a Kappa score > 0.3. This resulted in a set of non-redundant clusters, each represented by a statistically and semantically distinct term (Additional file 1: Table S7).

### Statistical analysis and visualization

For the COVID-19 dataset analysis, differential metabolites and proteins between outpatients and controls, as well as between severe/critical cases and controls were identified using linear models that adjusted for covariates, including sex, age, race, ethnicity and Charlson Comorbidity Index. To reduce confounding from external exposures such as pharmaceutical and dietary factors, xenobiotics were excluded from the differential metabolite analysis, thereby focusing on endogenous metabolic alterations. Significantly differential metabolites and proteins were identified using FDR-adjusted *p*-values < 0.05. Differential metabolites were used for ORA and TSA analysis. For MPNet module, differential metabolites and proteins for each comparison group were integrated as input. For MSEA analysis, the estimated coefficients for analyzed metabolites, derived from the linear models, were used as quantitative features to assess coordinated metabolic changes.

The R package clusterProfiler was used for KEGG and GO enrichment analysis of proteins [131]. The R packages pathview and SBGNview were used for the function module ORA and MSEA in the web server [132,133]; igraph (v2.0.3) and ggraph (v1.9.8) were used for metabolite-protein network visualization [128,134]; pheatmap (v1.0.12) was used for heatmap [135]; ggVennDiagram (v1.5.2) was used for Venn plot [136]. The other visualization was performed using ggplot2 (v3.5.1) [137].

## Declarations

### Ethics approval and consent to participate

Not applicable.

### Consent for publication

Not applicable

### Availability of Data and Materials

All data integrated into the HUBMet platform, including curated human blood metabolites, metabolite-protein associations, tissue relevance annotations, and analytical results, are freely accessible through the HUBMet web server at https://hubmet.app.bio-it.tech/home. The HUBMet web server front end is implemented using Vue 3, with backend data analysis scripts written in R and Python. The web server is deployed on Microsoft Azure. All data analysis scripts and associated data are publicly available on Zenodo: https://doi.org/10.5281/zenodo.17014155 [138].

## Authors’ contributions

W.Z. conceived and designed the study. X.W., X.Q., J.W., and W.Z. collected and contributed data to the study. X.W., X.Q., A.Z., W.Z. J.W., and S.B. performed the data analysis. W.Z., X.W., and X.Q. drafted the manuscript. All authors read and approved the final manuscript.

## Acknowledgements

The integrative multi-omics analysis of the processed data was performed using resources provided by SNIC through the Uppsala Multidisciplinary Center for Advanced Computational Science (UPPMAX) under Project Berzelius-2024-200. We thank the Beijing Cloudna Technology Co., Ltd., for technical support.

## Supplementary Information

Additional file 1: Supplementary tables

Additional file 2: Supplementary information and figures

